# Comparison and Optimization of Cellular Neighbor Preference Methods for Quantitative Tissue Analysis

**DOI:** 10.1101/2025.03.31.646289

**Authors:** Chiara Schiller, Miguel A. Ibarra-Arellano, Kresimir Bestak, Jovan Tanevski, Denis Schapiro

## Abstract

Studying the spatial distribution of cell types in tissues is essential for understanding their function in health and disease. A widely used spatial feature for quantifying tissue organization is the pairwise neighbor preference (NEP) of cell types, commonly referred to as co-occurrence or colocalization. Various methods to infer NEP have proved their utility in spatial omics studies, but despite their broad usage, no clear guidelines exist for selecting one method over the other. In this paper, we deconstruct frequently used methods into their underlying analysis steps and evaluate their optimal combination. We studied the methods on two aspects: (1) their discriminatory power to distinguish different tissue architectures and (2) their ability to recover the directionality of NEPs. We compared existing as well as our in-house developed method (conditional z-score (COZI)) and compared their performance using *in silico* tissue simulations and demonstrated its biological applicability in a myocardial infarction dataset. Overall, our study serves as a comprehensive guide for users and method developers in spatial omics analysis and offers a novel approach (COZI), which outperforms existing methods, to performing NEP analysis.

## Introduction

Understanding the spatial organization of tissues is crucial for identifying functional differences between conditions, such as healthy vs. diseased or respondent vs. non-respondent patients. In this context, spatial organization refers to the tissue architecture of cells and their interactions with each other. Cellular interactions and higher-order tissue architecture play a key role in determining tissue function, as they influence important processes such as immune responses and disease progression. With recent advances in spatial biology technologies^1–3^, researchers can now capture these spatial characteristics with unprecedented detail on a cellular and molecular level. Alongside technological progress, there is also a diversity of computational methods to analyze the spatial features of the resulting datasets^4,5^. One of the most widely used measures is the pairwise neighbor preference (NEP) of two cell types within tissues^6,7^. These methods, broadly used in spatial proteomics and transcriptomics, are commonly referred to as cell-cell interaction^8,9^, co-occurrence^10^, neighborhood enrichment^11^, or co-localization^12,13^ analysis methods. We will use the term neighbor preference (NEP) for a directional understanding of an index cell and its neighbors without ligand-receptor information.

The directionality of NEPs provides critical insights into tissue organization and functional dynamics. Some studies assume symmetry in their analysis, meaning that any given two cell types mutually prefer to be close to each other to the same extent^14,15^. Other studies provide NEP scores for two cell types in both directions, aiming to capture the directionality of NEPs^8,10,11^. Directional NEP can indicate patterns of communication or migration. For example, the spatial gradient of immune cells migrating from the stroma to tumor cores may indicate immune activation or evasion^16,17^. In wound healing, the state of infiltration of immune cells and their interaction with myofibroblasts corresponds to normal healing, cold or hot fibrosis^18,19^. Understanding and quantifying changes in these asymmetric spatial signatures is essential for comprehending tissue dynamics and developing therapeutic targeting strategies.

In general, NEP methods have provided new biological insights in various areas including oncology, immunology, and developmental biology. E.g., distinct NEPs of immune and tumor cells across patients in triple-negative breast cancer hinted towards general differences in immune cell infiltration in tumors and led to their sub-classification into cold, hot, and mixed tumors^10^. In another study, antigen-presenting cells near the epithelium or fibroblasts were more predictive for progressors than non-progressors in patients with invasive breast cancer^14^. A study focusing on glioblastoma compared to brain metastasis revealed a difference in cancer cells interacting with the bordering brain parenchyma^20^. Despite the power of these analyses, the methods used to infer NEPs vary greatly, with each approach offering distinct advantages and limitations.

This study explores the landscape of commonly used NEP methods, evaluating their performance on simulated and real-world datasets. We focused on the following methods providing NEP scores: histoCAT^8^, Spatial Enrichment Analysis (SEA)^10^, Giotto^15^, IMCRtools (classic)^21^, Squidpy^22^, Scimap^23^ and Misty^9^. We also included CellCharter^11^, which provides a function for NEP of niches, a function that can also be applied to cell phenotypes. There is no comparison or guide of these conceptually similar NEP methods and their underlying analysis steps to identify frequently occurring NEPs. Here, we deconstruct the methods into their underlying analysis steps and evaluate their performances on simulated ground-truth tissue data. We identify the ideal analysis steps for (1) their discriminatory power to distinguish different tissue architectures and (2) their ability to recover the directionality of NEPs.

Finally, we propose an approach that overcomes the limitations of existing NEP methods by integrating an optimal combination of the evaluated analysis steps. We identified that a conditional z-score (COZI) enables both the sensitive detection of differences in NEPs and their directionality. We demonstrate the biological utility of the method combination by applying it to a myocardial infarction dataset, where we analyze immune cell infiltration. Our findings highlight the heterogeneity of NEP method results, the advantages of combining the best analysis features from existing methods and offer a comprehensive guide for researchers and method developers in spatial omics analysis. By defining a common vocabulary, performing a systematic evaluation, and introducing a novel approach, we aim to enhance the accuracy and interpretability of spatial analyses.

## Results

### Unifying Concepts and Algorithms in NEP Analysis

To systematically compare existing pairwise neighbor preference (NEP) methods, we first identified their shared algorithmic components, key differences, and the terminology used to describe them. While these methods are widely applied in spatial biology, their underlying computational steps vary, making direct comparisons challenging. By deconstructing these methods into their core analysis steps, we established a unified framework that enables a structured evaluation of their performance (Fig. 1a-d). We identified three fundamental steps common to all NEP methods: (1) neighborhood definition, (2) neighbor quantification, and (3) NEP scoring (Fig. 1a-c). These steps and their combinations determine how spatial relationships between cell types are measured and how biological data can be interpreted. In the following, we describe each step and outline the algorithmic variations across methods.

**Fig. 1:**
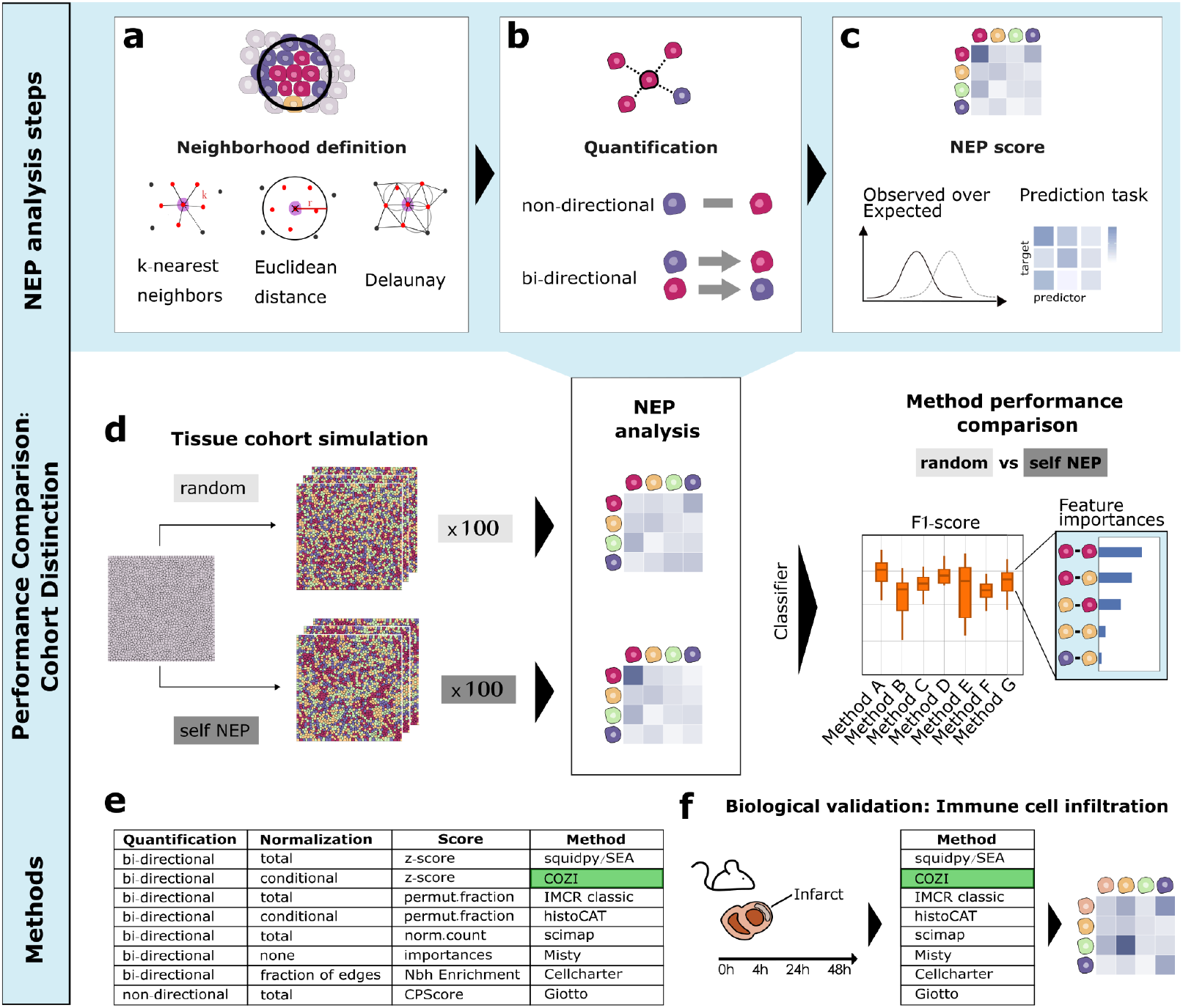
Overview of NEP method performance comparison. Three common NEP analysis steps were identified. **(a)** Neighborhood definition: Graph (Delaunay triangulation or k-nearest neighbors (kNN)) or distance based (Euclidean) neighborhood definition. User-provided parameters for Euclidean radius (r) or kNN (k) in red. **(b)** Quantification: Neighboring cells (within neighborhoods) are counted non- or bi-directionally. Non-directional counting aggregates neighbors (e.g., red-purple) without considering the index cell’s identity. Bi-directional counting assigns separate counts for each direction (e.g., “red-purple” vs. “purple-red”). **(c)** NEP score: NEP scores are determined. (Normalized) Neighbor counts are either assessed if they occur more often than expected or a prediction task is formulated for predictor-target cell types. **(d)** Schematic of the experimental design. Different NEP cohorts were designed and analyzed using the methods. A random forest classifier was trained with the NEP scores to evaluate how well the cohorts could be distinguished examining the F1 scores and feature importances. **(e)** Overview of methods and their underlying algorithmic steps to provide NEP scores. **(f)** Biological validation of the methods with a myocardial infarction dataset studying immune cell infiltration into the infarct core of mouse hearts.

#### Neighborhood definition

All methods first define a cellular neighborhood—the space around an index cell to determine its neighbors (Fig. 1a). This is based on either (i) a fixed distance (e.g., Euclidean in pixels/µm) or (ii) a graph-based approach (e.g., Delaunay triangulation, k-nearest neighbors). Most methods use a single neighborhood, except MISTy, which incorporates multiple distinct spatial views. We harmonized neighborhood definitions across methods for comparability of results.

#### Quantification

After defining the cellular neighborhood, the next step is quantifying neighboring cells within the specified space (Fig. 1b). This can be done in two ways: (i) Non-directional counting, which aggregates neighbors (e.g., red-purple) without considering the index cell’s identity, and (ii) bi-directional counting, which assigns separate counts for each direction (e.g., “red-purple” vs. “purple-red”). The latter can capture asymmetries in cell-cell preferences, revealing directional preference of one cell type to another. Most methods normalize neighbor counts using either “total” normalization, dividing by the total number of index cells, or “conditional” normalization, considering only index cells with at least one neighbor of the specified neighbor cell type. One exception: CellCharter does not normalize by the number of cells but rather the fraction of edges in the connectivity graph of an image.

#### NEP score

The final step is calculating a NEP score which is a unique score for each method (Fig. 1c). E.g., methods can either directly provide the scaled and normalized neighbor counts as NEP score, or assess and report whether a cell type pair is neighboring more often than expected by chance through permutation testing. Common statistical scores for NEP include z-scores, permutation fractions, and p-values. CellCharter reports the difference between the observed and the expected number of links derived from node degrees in the network, which is computationally faster than permutation testing^11^. MISTy treats NEPs as an inductive task, providing values for the variance explained^9^. Overall, all methods infer pairwise NEP scores, either non- or bidirectional.

Building on this systematic deconstruction of NEP analysis steps, we introduce the conditional z-score (COZI) as a novel approach not yet implemented by existing methods. Our evaluation of analysis steps, particularly those involving bi-directional counting and statistical scoring, showed that each analysis steps combination was implemented by a known method (Fig. 1e). However, one combination was missing: “conditional” normalization with a z-score, which no current method incorporates. To fill this gap, we developed and implemented COZI. In the following, we compare the performance of all existing methods and our in-house developed COZI (Fig. 1e) on simulated and biological data (Fig. 1f) and examine how differences in analysis steps explain performance variations and biological interpretability.

### Assessing Cohort Distinction Sensitivity of NEP Methods

We evaluated the methods by assessing their ability to distinguish tissue cohorts based on the spatial signatures they capture (Fig. 1d). A key goal in spatial biology is identifying distinctions between biological cohorts (e.g., healthy vs. diseased, responders vs. non-responders) based on tissue architecture. If a distinct NEP signature exists in one cohort but not the other, the methods should detect this difference. Therefore, instead of assessing how well methods reconstructed the underlying tissue structure, we initially focused on their ability to differentiate between the cohorts. Since real datasets lack ground truth spatial architecture, we used the *in silico tissue* (IST) generation framework^24^ to simulate tissue cohorts with known differences in NEP (Fig. 1d). To evaluate if the methods could capture the simulated cohort differences, we trained classifiers with each method’s NEP scores and evaluated the cohort classification performance by assessing F1-scores and feature importances (Fig. 1d). In the following, we systematically identify and explain performance differences based on the outlined NEP analysis steps.

### The Importance of Continuous Scores for Detection of NEP Differences

First, we simulated tissue cohorts for cohort distinction tasks, where the red cell type displayed varying levels of self-preference, ranging from random distribution to weak and strong self-preference (Fig. 2a). These different levels of self-preference were simulated across various abundances of the red cell type to evaluate the sensitivity of the methods for detecting NEPs of both low- and high-abundance cell types. We performed binary classifications comparing random vs. weak, random vs. strong, and weak vs. strong self-preference within abundance groups, ensuring that the comparisons focused on spatial patterns rather than differences in abundance. In the following analysis, we examine overall F1 scores, raw NEP scores, and F1 scores across increasing red cell type abundance, in order to assess general method performance and explain the observed performance differences (Fig. 2a).

**Fig. 2:**
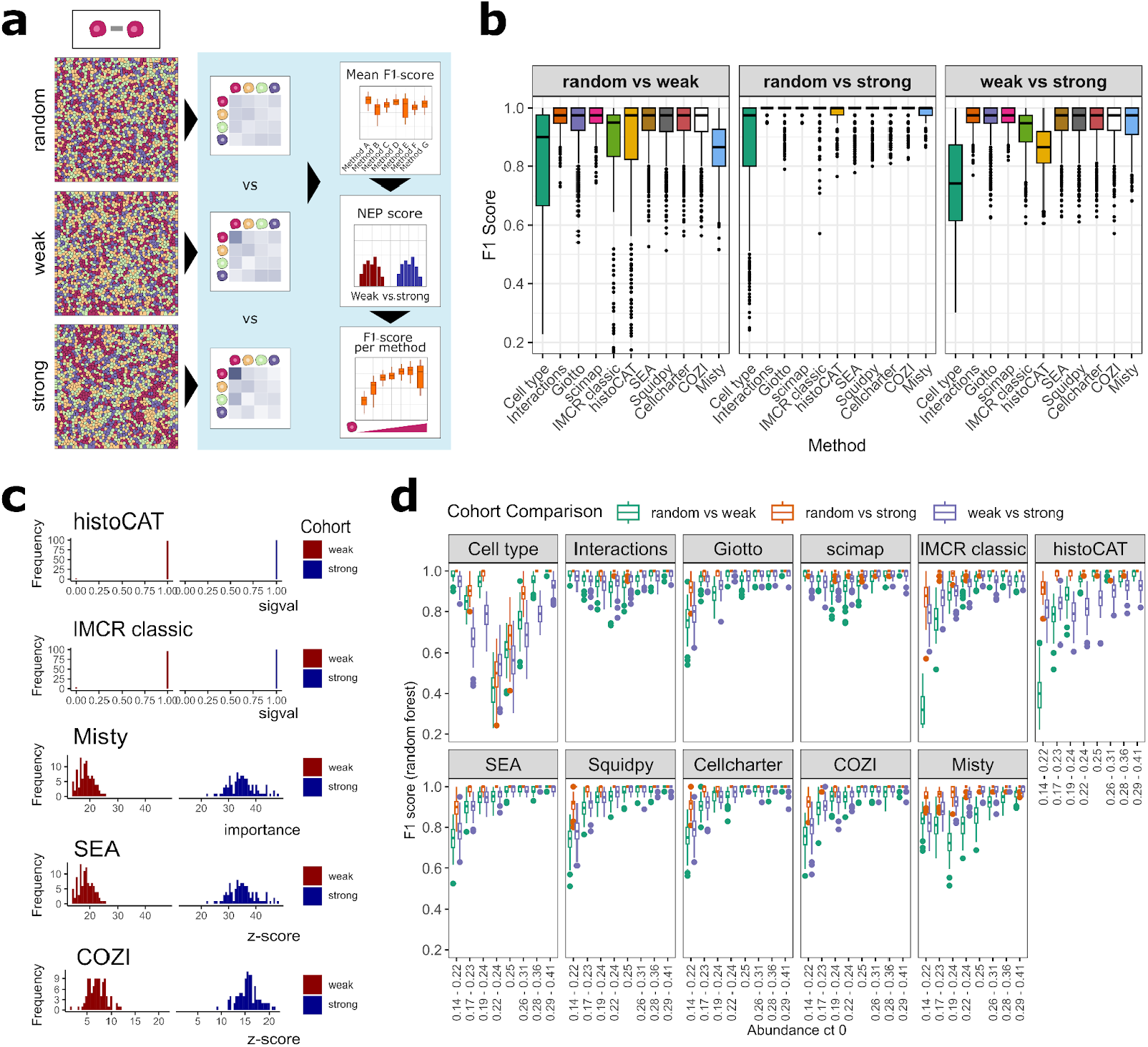
Systematic comparison of cohort distinction ability based on NEP analysis results with a random forest classifier. **(a)** Exemplary IST images for the three adjacency cohorts: random, weak, and strong self-preference of the red cell type. The red cell type was abundant at 25% in all depicted images. Schematic workflow: Cohorts were analysed by all NEP methods and binary classification was evaluated by assessing mean F1 scores (b), the true NEP scores (c) and the F1 scores across increasing abundance of the red (self-preference) cell type (d). **(b)** The boxplots show the mean of F1-scores per cohort distinction task across red cell type abundance groups. Cohort comparisons were made for random vs. weak, random vs. strong, and weak vs. strong self-preference. 100 images per cohort were simulated and analyzed. **(c)** NEP score distribution of red cell self interaction NEP scores of weak and strong self-preference cohorts from histoCAT, IMCRclassic, Misty, SEA, and COZI. All cell types were equally abundant at 25%. **(d)** The same experiment as in (b): F1 scores plotted per method across increasing cell type abundance cohorts of the red cell type.

All methods effectively distinguished the simulated tissue cohorts based on NEP differences, performing better than a random model and exceeding the classification based solely on cell type abundance differences (Fig. 2b, Supplementary Fig. 2a). The classification between the random and strong self-preference cohorts was nearly perfect across all methods (F1-score ∼1, Fig. 2b, second panel), as these cohorts exhibited the most pronounced spatial differences (Fig. 2a). Distinguishing between random and weak cohorts, and between the weak and strong cohorts resulted in slightly lower but still high performance (F1-score > 0.8, Fig. 2b, first and third panels). Of note, cohort classification based on cell type abundance differences alone also achieved F1-scores above 0.5 (random classification) due to inherent limitations in IST generation (Supplementary Fig. 1a and b). In this experimental setup, all methods performed very well and the only notable performance decline was observed for IMCRtools classic and histoCAT, particularly in the weak vs. strong cohort classification (Fig. 2c, third panel).

The performance drop of IMCRclassic and histoCAT can be attributed to their use of the permutation fraction metric for computing NEP scores. We examined the raw NEP scores of the red cell type to itself in weak and strong IST cohorts, each with 25% red cell type abundance (Fig. 2c). In both histoCAT and IMCRclassic, permutation fraction scores were largely 1, making it impossible to distinguish the red self-preference between the weak and strong cohorts. Both methods follow a simple question, assessing whether the underlying NEPs are significantly different from a random NEP. Therefore both methods provide a categorical response to a two-tailed test, with NEP scores of -1, 0, and 1. In cases of strong NEP, the score is 1 in most images, as nearly all randomly shuffled images have fewer neighbors than the original. In contrast, other methods like Misty (feature importances), SEA or COZI (z-scores) produce normally distributed NEP scores, allowing for clearer differentiation between weak and strong cohorts (Fig. 2c, Supplementary Fig. 3). NEP scores across all cell-cell interactions in the random, weak and strong cohort highlight only Scimap to not have normally distributed scores. Scimap scales NEP scores between -1 and 1, leading to skewed NEP scores even in the randomly organized tissue (Supplementary Fig. 3g).

We further evaluated the method performances across different red cell type abundance levels. The performance drop of IMCRclassic and histoCAT became more pronounced as the red cell type abundance increased (Fig. 2d). With higher abundance, methods with categorical NEP scores struggled to characterize subtle NEP differences, whereas methods with continuous scores performed better. Across different cell type abundance groups, Misty and Scimap exhibited F1 score trends similar to those driven by cell type abundance differences (Fig. 2d, first panel) or simple pairwise neighbor counts (Fig. 2d, second panel). Cohorts with the highest and lowest red cell type abundances were the easiest to distinguish based on simple cell type or neighbor counting (F1 ∼ 1). Misty and Scimap followed this trend, performing best when abundance differences were largest. In contrast, methods that use permutation testing or report differences between observed and expected NEPs performed worst at low red cell type abundances and best at high abundances with self-preference. This was expected, as NEPs of lowly abundant cell types are harder to capture compared to those of highly abundant cell types. Since Misty and Scimap do not compare observed to random NEPs, their NEP scores are more strongly influenced by cell type abundance differences. We will revisit this observation in our second simulation experiment.

### Unlocking NEP Directionality with Conditional Count Averaging

After evaluating the methods’ sensitivity to NEP differences, we investigated whether those providing bi-directional neighbor quantification (Fig. 1b) could capture NEP directionality using a simulated dataset. Here, directionality refers to one cell type showing a stronger preference for another as a neighbor than vice versa—for instance, immune cells infiltrating a tumor prefer tumor cells as neighbors more than the reverse. Therefore, we simulated a dataset with asymmetric cell-cell adjacencies, where the red cell type had a stronger preference for the yellow cell type than the other way around (Fig. 3a). We generated three levels of cross-preference — random, weak, and strong — while varying red cell type abundance. Although simulated and observed cell type abundances differed, we ensured cohorts included both higher red/lower yellow and lower red/higher yellow cell type abundances (Supplementary Fig. 1b). In this experiment, we added CellCharter’s package option to not include homotypic interactions, which we will call CellCharter* in the following.

**Fig. 3:**
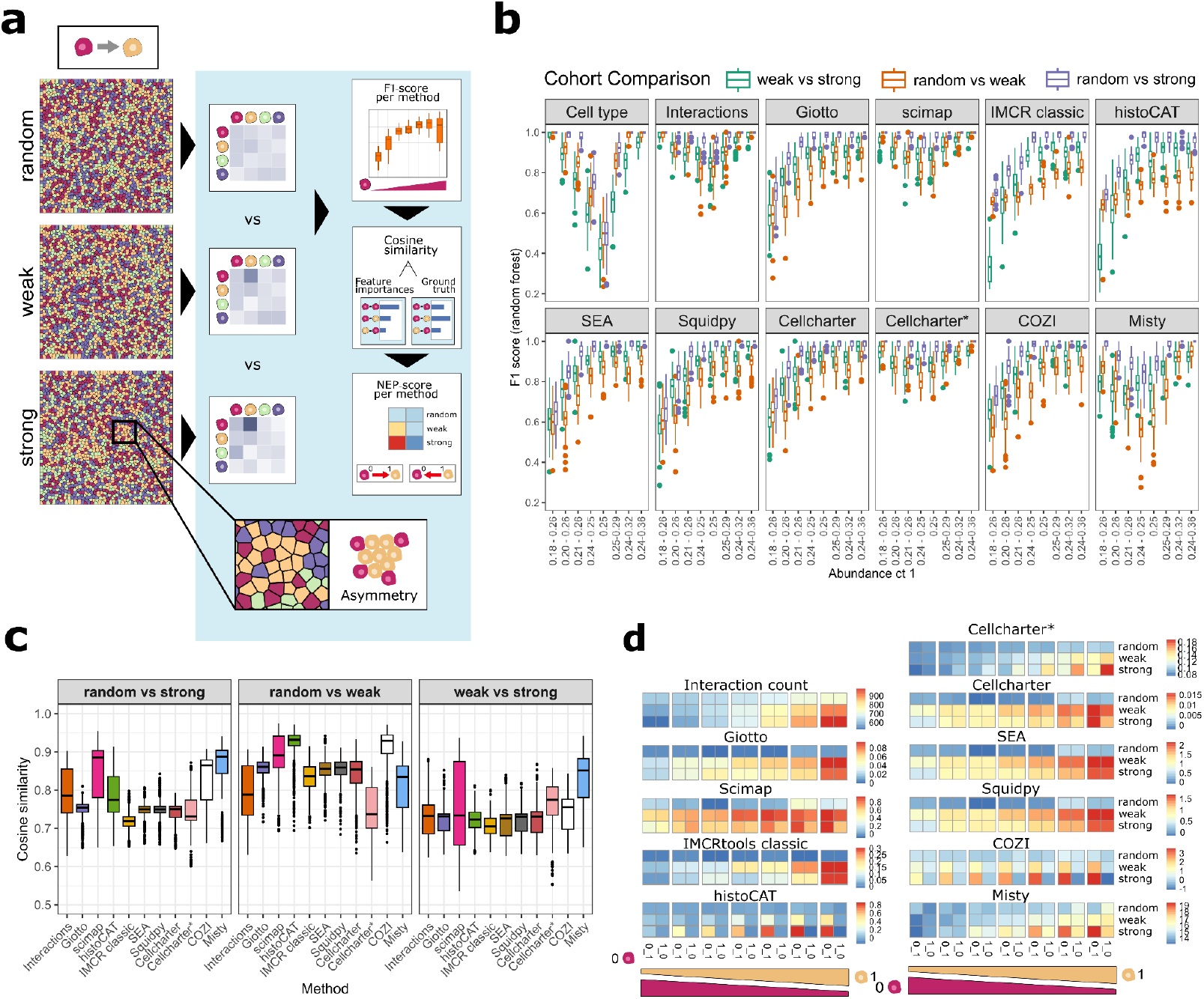
Systematic comparison of the ability to recover NEP directionality. **(a)** Exemplary IST images for the three adjacency cohorts - random, weak, and strong - with a red-to-yellow cell type cross-preference. The red cell type was abundant at 25% in all three depicted images. **(a)** Schematic depiction of (b)-(d). **(b)** F1 scores of cohort classification tasks per method across cohort comparisons. **(c)** Mean cosine similarities between feature importances and ground truth adjacency value differences between cohorts were calculated per method. Boxplots show mean cosine similarities per cohort distinction task. Cohort comparisons between random vs. weak, random vs. strong c and weak vs. strong cross-preference. **(d)** Raw NEP scores per method for the random, weak, and strong cohort for the red-yellow and the yellow-red NEP. Heatmap of interaction counts and NEP scores for Giotto, Scimap, IMCRtools classic, HistoCAT, CellCharter*, CellCharter, SEA, Squidpy, COZI, and Misty. Red cell type abundances range from higher than the yellow cell type to lower than the yellow cell type from left to right.

We first assessed cohort distinction performance using F1 scores. We found that all methods outperformed the cell type abundance and random label shuffling baselines, except for distinguishing random from weak self-preference, where the cell type abundance outperformed some methods (Supplementary Fig. 2b, Supplementary Fig. 4a). Across all methods, F1 scores increased with higher red cell type abundance (Fig. 3b). Misty and Scimap, as well as CellCharter*, followed similar F1-score patterns to the cell type abundance or interactions counts. For Giotto, we duplicated outputs to mimic bi-directional neighbor counting, enabling comparison of non-directional to bi-directional quantification. Giotto performed similarly well to SEA, COZI, CellCharter, and Squidpy in the cohort classification task — methods that inherently use bi-directional neighbor quantification (Supplementary Fig. 4a, Fig. 3b). Investigating the feature importances of the methods in the 25% abundance random vs. strong cohorts showed that only Misty, COZI and histoCAT had notably high feature importances for the red-yellow cross-preference (Supplementary Fig. 4b). CellCharter and Scimap showed high feature importances for the other direction, yellow-red, while none of the other methods captured a difference (Supplementary Fig. 4b). This led us to further investigate how and if these methods capture differences in directional NEP pairs.

While F1 scores measure overall classification performance in distinguishing different cohorts, cosine similarity specifically quantifies how well the methods recover the true adjacency patterns by comparing the methods-derived NEP scores to ground-truth cell-cell adjacencies. This distinction is crucial, as a method may achieve high classification performance without necessarily capturing the correct underlying spatial relationships. Therefore, we evaluated the recovery of differences in the ground truth cell-cell adjacencies by measuring cosine similarities between the model’s feature importances and ground-truth cell-cell adjacency values (Fig. 3c). Higher cosine similarity indicates better alignment with the true adjacency structure. A key difference emerged between neighbor count averaging methods: The bi-directional methods SEA, IMCRtools classic, CellCharter and Squidpy yielded cosine similarities comparable to Giotto, the only non-directional method (Fig. 3c). The three methods — SEA, IMCRtools classic, and Squidpy — normalize neighbor counts by the total number of cells of the same type as the index cell. Their raw NEP scores for red-yellow and yellow-red were indistinguishable, confirming that the methods do not capture NEP directionality (Fig. 3d,e). In contrast, histoCAT and COZI showed higher cosine similarities than the named bi-directional methods (Fig. 3c). Both perform conditional normalization, normalizing index-neighbor counts by the number of cells of the index cell type with at least one link to a neighbor cell type. This approach leads to the recovery of the simulated red-yellow NEP but not vice versa consistently across abundance levels (Fig. 3d). Both versions of CellCharter also capture directionality, as stated in the method. CellCharter including homotypic interactions captures a correct directional red-yellow preference in cohorts with higher abundance of the yellow cell type, while it does not capture directionality in equal or low red abundance cohorts (Fig. 3d). Without permutation testing in CellCharter, there is a potential cell type abundance bias. The total number of cells of the red type is proportional to the total number of edges which decreases the ability to capture the preference of a highly to a lowly abundant cell type. CellCharter* excluding homotypic edges in the network consistently captures NEP differences between the two directions across cell type abundance differences, but the other way around than simulated (Fig. 3d).

Misty and Scimap also reached high F1-scores (Supplementary Fig. 4a), cosine similarities (Fig. 3c), and differences between the red-yellow and yellow-red neighbor preferences (Fig. 3d). Both methods, however, were also affected by cell type abundances throughout this study (Fig. 3b). Both methods falsely recover a higher score for the yellow-red than for the red-yellow NEP for low abundances of the yellow cell type (Fig. 3d). This observation is explainable by their score: Scimap does not provide a z-score, but the scaled neighbor count of the red-yellow cell co-occurrences divided by the number of red cells. A low number of yellow cells inflates the yellow-red score compared to red-yellow. Similarly, Misty provides values of the variance explained based on a prediction task not corrected by permutation testing. As Scimap and Misty do not provide scores based on permutation testing or expected neighbor counts, both do not account for differences in cell type abundances — in comparison to how a permutation test does. Therefore, it is questionable how well they can correctly recover NEPs of a highly abundant cell type to a lowly abundant one.

Overall, conditional averaging was the decisive algorithmic step in recovering NEP directionality in our simulation experiments, making it a key feature in the discussed methods.

The combination of conditional count averaging and z-score calculation performed best across all simulation experiments in this study. Due to its limitation to distinguish similar NEPs with varying degrees of strength, histoCAT’s distinction ability drops when comparing F1 scores of the weak vs the random cross-preference cohort, thus confirming our previous results (Supplementary Fig. 4a). The COZI performance stays as high as the other permutation-based methods and recovers directionality. Giotto, IMCRtools classic, SEA, and Squidpy do not recover directionality due to total count normalization. CellCharter and CellCharter* do capture directionality but either not across all cell type abundance groups or in the opposite direction. Scimap and Misty are primarily affected by cell type abundance differences.

In summary, we recommend a combination of advantageous algorithmic features to have a sensitive and directional NEP score with COZI. In the following, we will apply the methods to a biological dataset to illustrate how their differences impact biological interpretability.

### Recovery of Tissue Structures in a Myocardial Infarction Dataset

We applied COZI as well as the other methods to a myocardial infarction dataset from Wuennemann *et al*., which studied immune cell infiltration into the mouse heart infarct area at multiple time points after myocardial infarction (4, 24, and 48 hours)^25^. At all time points, the injured region containing stressed (Ankrd1+) cardiomyocytes is visible (red, Fig. 4a). Both visual inspection (Fig. 4a) and cell count analysis of manually annotated regions (Fig. 4b) confirm that neutrophils infiltrate the injury earlier than Ccr2+ monocytes/macrophages, referred to as monocytes from here on. The original study reported that immune cell infiltration, particularly of monocytes, in the acute phase occurs through the endocardium—the inner layer of the heart—specifically in the left ventricle (Fig. 4b, d). These observations serve as biological anchor points to evaluate the performance of the different methods. We systematically examined the visual spatial distributions, region-specific cell type abundances, and the computed NEP scores to guide biologically meaningful interpretations that can be applied to other datasets.

**Fig. 4:**
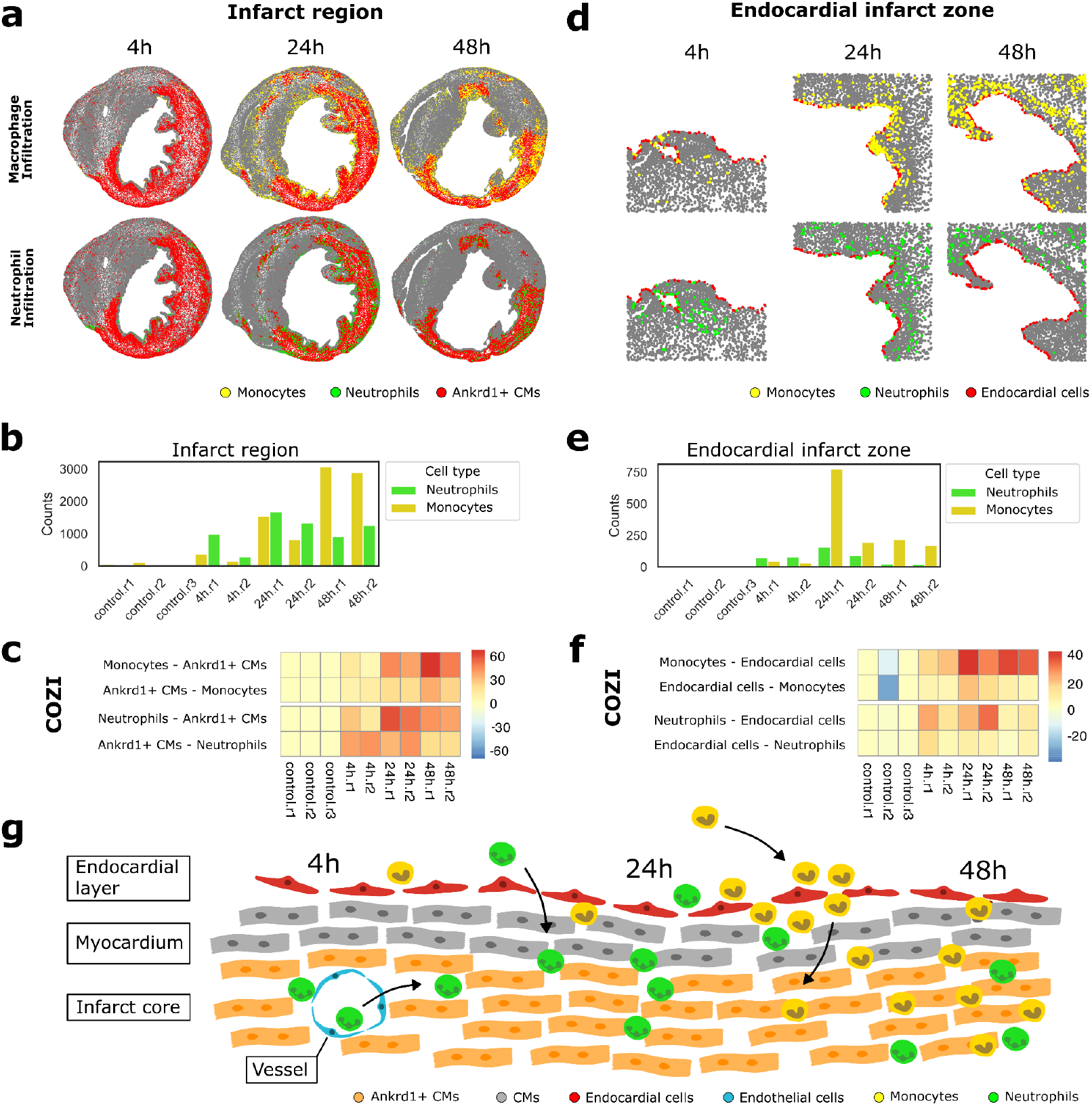
Recovery of tissue structures in a myocardial infarction dataset. **(a)-(b)** Monocyte (yellow) and neutrophil (green) infiltration into the (a) infarct region (red) (4h.r2, 24h.r1, 48h.r1) and through the (b) endocardial layer (red) (4h.r1, 24h.r1, 48h.r1) at 4, 24 and 48h after infarct. Scatter plot colored by cell type identities. **(c)-(d)** Cell type percentages of monocytes and neutrophils in the (c) infarct region and the (d) endocardial infarct zone. **(e)-(f)** COZI neutrophil and monocyte NEP scores with (e) Ankrd1+ cells and (f) endocardial cells for control, 4, 24, and 48h samples. **(g)** Schematic of biological findings for immune cell infiltration through the endocardium or blood vessels into the injury site along the timeline after infarction from left to right. Endocardial cells (red), myocardium (grey), Ankrd1+ stressed cardiomyocytes (orange), endothelial cells (blue), neutrophils (green), and monocytes (yellow).

We first investigated how well the methods captured immune cell infiltration into the infarct and showed that only COZI successfully identified the earlier infiltration of neutrophils compared to monocytes while preserving directionality in NEP (Fig. 5c, Supplementary Fig. 5).

#### Monocyte infarction infiltration

Monocytes show almost no infiltration into the infarct at 4h (Fig. 4a, b). At 24 hours, monocytes start infiltrating (Fig. 4a, b) and COZI shows a strong preference of monocytes for Ankrd1+ cardiomyocytes (z-score ∼40, Fig. 4c), and the reverse preference is absent. This observation suggests that monocytes are infiltrating the injury site by surrounding the infarct, while Ankrd1+ cells remain clustered together—similar to the asymmetric pattern observed in our simulations, where yellow cells clustered together and red cells surrounded them (Fig. 3a). By 48 hours, monocytes are fully integrated into the infarct, leading to more similar NEP scores between the two cell types (z-score ∼30-40, Fig. 4c). CellCharter* and Scimap consistently capture the peak preference at 48h as well, while they do not detect an infiltration at 24h. CellCharter, Scimap, SEA and Squidpy all capture an avoidance in samples 4h.r2 and 24h.r1. At 4h.r1, there are almost no monocytes (Supplementary Fig. 8b) while they visually do not seem to avoid the infarct region (Supplementary Fig. 6b). At 24h, monocytes are surrounding the infarct core visibly (Figure 4a). Misty, histoCAT and COZI, the three methods which are capturing directionality best in the simulation experiment, detect a monocyte preference for Ankrd1+ cardiomyocytes at 24h. In contrast, CellCharter, Scimap, SEA and Squidpy (Supplementary Fig. 5) report an avoidance of monocytes for Ankrd1+ cells for this time point.

#### Neutrophil infarction infiltration

Neutrophils already start infiltrating the infarct region at 4h (Fig. 4a, c, Supplementary Fig. 6d). Looking at COZI results, Ankrd1+ cardiomyocytes show a preference for neutrophils already at 4 hours (Fig. 4c). Interestingly, the preference pattern is reversed compared to expectations during the infiltration phase observed with monocyte preference for Ankrd1+ cells, prompting further investigation of the infiltration pattern. In 4h.r2, neutrophils seem to form clusters of cells within the infarct region being surrounded by Ankrd1+ cells (Fig. 4a, lower panel). They are more abundant and mixed in 24h.r1 leading to mutual NEPs between neutrophils and Ankrd1+ cells (Fig. 4e, Supplementary Fig. 6d). We found that neutrophils primarily infiltrate through blood vessels within the infarct rather than from the surrounding tissue (Supplementary Fig. 8a). This was validated by a preference of endothelial cells (which form vessel structures) for neutrophils at 4 hours — a pattern again only recovered by COZI (Supplementary Fig. 7e). By 24 hours, neutrophils are fully present within the Ankrd1+ infarct core, leading to mutual preference between the two cell types. At 48 hours, neutrophils show a preference for Ankrd1+ cells, as they become fully embedded in the infarct, while the Ankrd1+ cells gradually die off, leaving remaining cardiomyocytes surrounded by neutrophils. All other methods capture a neutrophil infiltration rather than a monocyte infiltration, while only some capture a consistent NEP already at 4h. None of the other methods capture a change in directionality along the time axis.

Immune cell infiltration through the endocardium was a key finding of the myocardial infarction study, particularly for monocytes. We could verify this directional NEP only with COZI.

#### Monocyte endocardium infiltration

We examined NEP scores between endocardial and immune cells over time after the infarct. The images clearly show starting infiltration at 4h, a strong infiltration at 24h and further infiltrated monocytes at 48h (Fig. 4b). Looking at COZI scores at 4h, monocytes have a slight preference for endocardial cells, while at 24 hours, monocytes show a strong preference for endocardial cells (z-score ∼40), and endocardial cells also exhibit an elevated preference for monocytes (z-score ∼20), reflecting the peak of infiltration through the endocardium (Fig. 4d, f). At 48 hours, monocytes are observed deeper within the tissue, reducing neighbor counts with endocardial cells. Endocardial cells lose their NEP for monocytes as large areas of the endocardial layer are no longer neighboring monocytes, while monocytes are migrating. All other methods capture a peak infiltration through the endocardium at 24h, while most are largely resembling a strong increase in monocyte abundance and interaction count in sample 24h.r1 (Supplementary Fig. 9). Additionally, CellCharter, Misty, and Scimap report a preference of endocardial cells for monocytes, in contrast to COZI, which shows the opposite direction (Supplementary Fig. 9). The lower abundance of endocardial cells compared to monocytes at 24h highlights the method’s limitation in capturing preference from highly abundant to lowly abundant cell types (Supplementary Fig. 8b).

#### Neutrophil endocardium infiltration

Neutrophils show a weaker preference for endocardial cells at 4 hours (z-score ∼10-20) than monocytes, indicating less infiltration through the endocardium than monocytes. By 24 hours, neutrophil infiltration through the endocardium increases but remains weaker than monocyte infiltration, with endocardial cells not exhibiting the same strong preference for neutrophils. This is reflected in the images and the decline in neutrophil infiltration by 48 hours (Fig. 4b, d). Notably, COZI was the only method that captured neutrophil infiltration through the endocardium at a biologically meaningful level, both overall and at specific time points (Fig. 4f, Supplementary Fig. 9).

## Discussion

This study provides a comprehensive guide over NEP analysis methods and proposes an optimal combination of analysis steps for gaining biologically meaningful insights. This work highlights the heterogeneity of the available methods and their results, and is a valuable resource for researchers, helping them make informed decisions for their spatial analysis. Additionally, we demonstrate how IST-generated spatial data can effectively be used to compare spatial omics methods.

When evaluating NEP methods, we found that all methods could distinguish between tissue cohorts based on spatial differences. However, minor algorithmic differences significantly impacted the ability to recover biologically relevant spatial patterns and led to vastly heterogeneous results. SEA, Giotto, IMCR classic, and Squidpy did not recover NEP directionality, a crucial aspect of spatial organization. For example, during immune cell infiltration into the tissue, the infiltrating cells initially prefer resident tissue cells, but once fully integrated, both cell types exhibit mutual preferences for each other. Moreover, tissue architecture is inherently asymmetric, meaning that cell-cell adjacency relationships should not be assumed to be bidirectional. Methods that fail to capture directionality lose this critical information.

To address this, we demonstrated that conditional averaging is essential for recovering meaningful directionality by normalizing interaction counts exclusively among interacting cells. Our results showed that COZI successfully captured directional relationships in simulated and biological datasets. While CellCharter, Misty, and Scimap also recovered directionalities in their interaction scores, the methods were sensitive to differences in cell type abundances. Specifically, they could only detect preferences from a lowly abundant cell type toward a highly abundant one, but not vice versa. It is worth noting that Misty was not primarily designed for cell type level NEP analysis but can be adapted for this purpose by bypassing intraview calculations.

Since this is the first study to perform a method comparison using the IST framework from Baker *et al*.^14^, we also identified some limitations associated with this approach. One key limitation was simultaneously maintaining predefined cell type abundances and adjacency constraints. When a low-abundance cell type was required to maintain high adjacency with multiple cell types, there were often too few cells to comply with both conditions. During these simulations, the IST framework prioritizes adherence to the cell-cell adjacency parameter over maintaining exact cell type abundances. As a result, cell-cell adjacencies were reliably simulated while deviations from the given cell type abundances occurred. Understanding these inherent tissue architecture limitations – e.g. that a low abundant cell type cannot have very high preferences for all other cell types – can help setting realistic expectations when analyzing biological tissues. Despite these challenges, IST-generated data allowed us to identify algorithmic differences between methods that might have been overlooked in real-world datasets. One could also expand the simulation framework to 3D simulations, addressing the increasing availability of 3D spatial omics datasets.

NEP analysis can also be performed and assessed on 3D datasets. E.g., COZI can be run on 3D datasets. However, the COZI score is affected by the total number of cells, which is much more variable in 3D datasets than in 2D datasets. The higher the cell number, the higher the COZI score due to the effect size of z-scores. We therefore recommend dividing COZI scores of a sample by the square root of the number of cells in that sample when encountering large differences in cell type numbers.

Expanding NEP analysis beyond local cell-cell neighbors could provide deeper insights into global tissue architecture. A promising approach is the combination of a conditional z-score with a multiview framework like Misty, which captures multiple spatial neighborhood views at once. While NEP methods like COZI can define larger neighborhoods, the defined space includes direct and distant neighbors at the same time. In contrast, Misty and other multiview approaches^4^ allow for the independent assessment of short- and long-range spatial neighbors. Future work could explore how to best integrate these approaches for a more comprehensive understanding of global tissue organization.

The IST framework used in this study was designed to simulate local tissue architecture, which aligns with our focus on NEP analysis at the local level. However, biological tissues are organized into more complex structures, such as cellular niches, gradients, and functional domains. Expanding the analysis beyond NEP to identify spatially coherent multicellular structures will provide a more comprehensive understanding of tissue function and disease progression. Nevertheless, statistical scores like COZI are both easy to interpret and robust, making them suitable for future applications.

## Methods

### Tissue data generation

#### *In silico* tissue generation framework

We generated *in silico* tissues (IST) using the open-source IST generation method from Baker *et al*. 2023^24^. We used the provided Python scripts for tissue scaffold generation and cell type label assignment (klarman-cell-observatory/PowerAnalysisForSpatialOmics).

First, blank tissue scaffolds were generated with a random-circle packing algorithm. Parameters for the field of view (FOV) size *f* and r_max_ and r_min_ radius of cells were varied. In the final dataset, we used *f* = 1000, r_min_ = 10, and r_max_ = 10. The circle representation was converted into a Voronoi graph representation for downstream cell type annotation. The script was adapted to output the tissue matrices of the scaffolds for further annotation. The nodes of the unlabeled tissue scaffold were annotated using the heuristic approach due to speed and practicability. A vector *p* containing the abundances for each of the *k* cell types was specified.

A *k* x *k* matrix *H*_*k* x *k*_ defines the probability that a cell *k*_*i*_ is adjacent to a cell *k*_*j*_. In the original method’s use case, *p* and *H*_*k* x *k*_ were estimated from the provided biological tissue. Here, we provided the parameters to generate tissue architecture cohorts with similar cell type abundances. The image was partitioned into a grid of regions with size t x t pixels for heuristic annotation. We used a grid size of 200. Within an initial random region of blank nodes, the cell type labels were sampled from a multinomial distribution, with *p*_*k*_ being the cell type distribution. The neighborhood graph distance was set to 2. Depending on *k*, the neighboring node identities were sampled from a multinomial distribution of the respective row vector of the adjacency matrix *H*_*ki* x *k*_. The region of the grid partition was shifted by *t/2* in the x and y direction, and the annotation was repeated for every unlabeled node. After completion, random nodes were selected, and the neighborhood composition was calculated and compared with *H*_*k* x *k*_. Nodes with overabundant cell type labels were swapped to under-abundant ones. The number of final iterations *I* was 300 and cell type label swaps *S* was 50.

#### Dataset and cohort design

We simulated cohorts with differing cell-cell adjacencies and cell type abundances of 4 cell types (Table 1). Across cohorts, we used the same tissue scaffolds for cell type annotations. We adapted *H*_*k* x *k*_ and *p* to generate one symmetric dataset D1 (Table 2), and an asymmetric dataset, D2 (Table 3). Per dataset, we generated three tissue architecture levels for cohort generation. We simulated D1 with a random (H_0×0_ = 0.25), a weak (H_0×0_ = 0.45), and a strong (H_0×0_ = 0.6) self-preference of cell types 0 (Table 2). For D2, we simulated a random (H_0×1_ = 0.25), a weak (H_0×1_ = 0.45), and a strong (H_0×1_ = 0.6) cross-preference between cell type 0 and 1 (Table 3). Per adjacency level, the abundance of cell type 0 *p*_*k=0*_ ranged from 0.05 to 0.55, with p_k={1, 2, 3}_=(1-*p*_*k=0*_)/3 (Table 1). Per dataset (D1 and D2), we combined the eight vectors for *p* with the three different adjacency matrices H_*k* x *k*_ for cell type label assignment. Therefore, we generated 24 different cohorts with 100 samples each per dataset.

**Table 1:** Cell type abundance vectors p_k1-4_ for ranging cell type abundances of cell type 0 for D1 and D2.

| Cell type | ab0_0.05 | ab0_0.1 | ab0_0.15 | ab0_0.2 | ab0_0.25 | ab0_0.35 | ab0_0.45 | ab0_0.55 |
| --- | --- | --- | --- | --- | --- | --- | --- | --- |
| $k_0$ | 0.05 | 0.1 | 0.15 | 0.2 | 0.25 | 0.35 | 0.45 | 0.55 |
| $k_1$ | 0.32 | 0.3 | 0.28 | 0.27 | 0.25 | 0.22 | 0.18 | 0.15 |
| $k_2$ | 0.32 | 0.3 | 0.28 | 0.27 | 0.25 | 0.22 | 0.18 | 0.15 |
| $k_3$ | 0.31 | 0.3 | 0.29 | 0.26 | 0.25 | 0.21 | 0.19 | 0.15 |

**Table 2:** Adjacency matrices H_kxk_ for investigating self-preference of cell type 0 in D1 on three different levels.

| A: Random |  |  |  |  | B: Weak self preference 0-0 |  |  |  |  | C: Strong self preference 0-0 |  |  |  |  |
| --- | --- | --- | --- | --- | --- | --- | --- | --- | --- | --- | --- | --- | --- | --- |
| | $k_0$ | $k_1$ | $k_2$ | $k_3$ | | $k_0$ | $k_1$ | $k_2$ | $k_3$ | | $k_0$ | $k_1$ | $k_2$ | $k_3$ |
| $k_0$ | 0.25 | 0.25 | 0.25 | 0.25 | $k_0$ | 0.45 | 0.18 | 0.18 | 0.19 | $k_0$ | 0.60 | 0.13 | 0.13 | 0.14 |
| $k_1$ | 0.25 | 0.25 | 0.25 | 0.25 | $k_1$ | 0.18 | 0.27 | 0.28 | 0.27 | $k_1$ | 0.13 | 0.29 | 0.29 | 0.29 |
| $k_2$ | 0.25 | 0.25 | 0.25 | 0.25 | $k_2$ | 0.18 | 0.28 | 0.27 | 0.27 | $k_2$ | 0.13 | 0.29 | 0.29 | 0.29 |
| $k_3$ | 0.25 | 0.25 | 0.25 | 0.25 | $k_3$ | 0.19 | 0.27 | 0.27 | 0.27 | $k_3$ | 0.14 | 0.29 | 0.29 | 0.28 |

**Table 3:** Adjacency matrices H_kxk_ for investigating cross-preference of cell type 0 to cell type 1 in D2 on three different levels.

| A: Random |  |  |  |  | B: Weak cross preference 0-1 |  |  |  |  | C: Strong cross preference 0-1 |  |  |  |  |
| --- | --- | --- | --- | --- | --- | --- | --- | --- | --- | --- | --- | --- | --- | --- |
| | $k_0$ | $k_1$ | $k_2$ | $k_3$ | | $k_0$ | $k_1$ | $k_2$ | $k_3$ | | $k_0$ | $k_1$ | $k_2$ | $k_3$ |
| $k_0$ | 0.25 | 0.25 | 0.25 | 0.25 | $k_0$ | 0.18 | 0.45 | 0.18 | 0.19 | $k_0$ | 0.13 | 0.60 | 0.13 | 0.14 |
| $k_1$ | 0.25 | 0.25 | 0.25 | 0.25 | $k_1$ | 0.27 | 0.18 | 0.28 | 0.27 | $k_1$ | 0.29 | 0.13 | 0.29 | 0.29 |
| $k_2$ | 0.25 | 0.25 | 0.25 | 0.25 | $k_2$ | 0.28 | 0.18 | 0.27 | 0.27 | $k_2$ | 0.29 | 0.13 | 0.29 | 0.29 |
| $k_3$ | 0.25 | 0.25 | 0.25 | 0.25 | $k_3$ | 0.27 | 0.19 | 0.27 | 0.27 | $k_3$ | 0.29 | 0.14 | 0.29 | 0.28 |

Of note, IST generation has local tissue organization constraints and cannot build global relationships in a sample. However, the NEP methods we compared are restricted to local tissue organization and thus are well suited to this setup. We further performed quality control of the simulated data and identified discrepancies between the simulated and the observed cell type abundance levels (Supplementary Fig. 1a and b, Methods). This discrepancy was most apparent in cohorts with lowly abundant red cell type or very highly abundant red cell type but not apparent in cohorts with similar cell type abundances across cell types. Therefore, our study revealed critical limitations in the published framework.

### Neighbor Preference (NEP) Methods

#### histoCAT and IMCRtools

The IMCRtools R toolbox provides a classic and histoCAT version to find statistically enriched NEPs of cell types in defined cellular neighborhoods (https://github.com/BodenmillerGroup/imcRtools). The original neighborhood definition of histoCAT is distance-based^8^. We performed the classic and histoCAT analysis of simulated data with a Delaunay graph neighborhood definition, as provided in IMCRtools for comparability. For the biological dataset we used k=5. NEPs are counted bi-directionally, so the method provides preference values for cell type A to type B and from type B to type A. The neighbor count is averaged in two ways: the classic method divides the interaction count between A and B by the total number of cells of type A in the sample, while HistoCAT divides the interactions between A and B by the number of type A cells with at least one neighbor of type B^8^. We termed this “conditional averaging” throughout the manuscript. After counting, the cell type labels are randomly permuted (*n* = 300 times), and randomized interactions are counted and averaged as described. If an interaction is abundant significantly more or less is indicated in the NEP score “sigval” with 1 and -1 respectively. Not significant interactions are reported as 0. We obtained a sample-by-interaction matrix with sigval values as output for downstream evaluation.

#### Giotto

The Giotto NEP analysis was performed with the Giotto R package. The method defines neighbors by Delaunay triangulation. Giotto counts interactions non-directionally, providing only one value for the interaction A-B. We permuted cell type labels randomly *n* = 300 times, forming a null distribution. The true interaction count is divided by the mean null distribution count and termed cell proximity score (CPscore). A wrapper of the function is written in cellProximityEnrichment() of the package. We obtained a sample-by-interaction matrix with CPscores as output for downstream evaluation.

#### Spatial enrichment analysis (SEA)

Spatial enrichment analysis (SEA) detects statistically enriched cell-cell interactions^10^. The method is described and used in several papers^10,14^ but not provided as a stand-alone package. The authors kindly shared Matlab and Python code for analysis, which we adapted on a fork of Scimap to use their underlying neighborhood definition and permutation framework. The method was initially described using an Euclidean distance neighborhood definition. We implemented the method with a bi-directional count, while non- and bi-directional counts were both described in studies performing SEA. We ran the method with a Delaunay graph neighborhood definition for the simulated and knn (k=5) for the biological dataset and permuted the cell type labels *n* = 300 times. The z-score is calculated as:

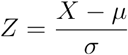

Where:

*X* = observed neighbor count

*μ* = mean of null distribution of X in permutation

*σ* = standard deviation of null distribution of X in permutation

We obtained a sample-by-interaction matrix with z-scores as output for downstream evaluation.

#### MISTy

The Multiview Intercellular SpaTial modeling framework (MISTy)^9^ enables analysis of the relationships of markers or cell types in spatially resolved data. The machine learning framework allows flexible definitions of spatial contexts (views). These views can, for example, be the inside of a cell (intraview), the direct surrounding of a cell (juxtaview), or a larger environment around a cell (paraview). MISTy models the complete tissue interactions for predictor/target relationships in each view. Each view-specific model quantifies the variance explained by each marker within that view for predicting the presence of the target cell type or marker. MISTy is implemented in R (Version 1.6.1.), and we adapted the script for Fig. 2 in the original publication for our analysis9,26 (https://github.com/saezlab/misty_pipelines). We used MISTy with cell type annotations only (one-hot encoded cell types), which bypasses the intraview modeling. We defined the immediate neighborhood as juxtaview with a neighborhood threshold of 40 for Delaunay triangulation for the simulated and with knn (k=5) for the biological dataset. We ran MISTy on the defined juxtaview with the run_misty() function, setting the argument bypass.intra to TRUE. We did not summarize the results but performed a sample-wise analysis. We obtained a sample-by-predictor/target matrix with model variances explained as NEP scores as output for downstream evaluation.

#### Squidpy

The spatial analysis suite Squidpy provides a nhood_enrichment function for calculating NEPs for cell types in tissues. It follows the same approach as SEA and randomly permutes cell type labels to generate a null distribution for calculating z-scores. It divides the counted interactions by the total number of cells and generates a symmetric NEP score. Squidpy outputs the counted interactions, which we used for our interaction count comparison, as well as the calculated z-scores, which we used as NEP from Squidpy. We ran the method with a Delaunay graph neighborhood definition for the simulated and knn (k=5) for the biological dataset and permuted the cell type labels *n* = 300 times. We obtained a sample-by-z-score matrix as output for downstream evaluation of Squidpy and a sample-by-count matrix as output for downstream evaluation of the interaction counts.

#### Scimap

The spatial analysis suite Scimap provides a spatial_interaction() function for calculating NEPs of cell types in tissues. The method also performs permutation tests to infer significant neighbor preferences in the tissue. While the method calculates p-values based on absolute difference or z-score-based significance, the output score is not the calculated z-score. The Scimap interaction score is the cell type abundance normalized and scaled number of interactions in the tissue. Therefore, the score does not include information from the permutation test, but significant interactions can be filtered based on the provided p-values. The original function only allows for the definition of k-nearest neighbors or Euclidean distance-based neighborhoods. We implemented a Delaunay graph neighborhood definition to compare it to the other methods. We ran the method with a Delaunay graph neighborhood definition for the simulated and knn (k=5) for the biological dataset and permuted the cell type labels *n* = 300 times. We obtained a sample-by-score matrix as output for downstream evaluation.

#### CellCharter

CellCharter is a spatial analysis suite and offers an nhood_enrichment() function for computing NEPs between clusters in a spatial graph. In the publication, this function is proposed for NEPs between cells of niches, while we ran the function for NEPs between cells of phenotypes without changing anything in the function. The method does not perform permutation testing but, in the asymmetric version that we used, calculates the difference between the observed and the expected neighbor edges in an analytical formula. We ran the method with only_inter=False as CellCharter and only_inter=True as CellCharter* to either include or exclude links between homotypic clusters, respectively^11^. We ran the method with Delaunay 1-hop neighborhood definition for the simulated and the biological dataset. We obtained a sample-by-score matrix as output for downstream evaluation.

#### Conditional z-score (COZI)

The conditional z-score (COZI) outputs normalized z-scores as NEP scores. Let *C*_*i*_ and *C*_*j*_ be subsets of *C*, where each *C*_*k*_ represents all cells of a specific cell type *k* in the image. The boolean adjacency matrix *A* contains all neighboring cell edges, where *A*_*st*_ indicates the presence of an edge between cells s and t.

The observed number of edges between *C*_*i*_ and *C*_*j*_ is given by:

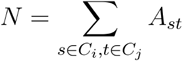

To account for cell abundances, we normalize the observed interactions *N* by *M*, the number of cells of type *C*_*i*_ with at least one edge to a cell of type *C*_*j*_:

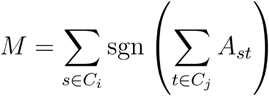

This results in the normalized interaction count *O*.

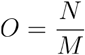

To assess the significance of observed interactions, COZI performs permutation testing by randomly shuffling the cell type labels across cell locations. In each permutation, neighbor counts are recomputed using the same normalization procedure to obtain a null distribution for *O*.

The final COZI score is the z-score, of the observed *O* relative to its permuted distribution, using the mean *μ*_*perm*_ and standard deviation *σ*_*perm*_ of *O*_*perm*_ across permutations.

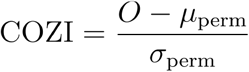

A high COZI score indicates stronger-than-random spatial association, while a low or negative score suggests no significant spatial preference or avoidance. COZI is affected by differences in total cell type numbers. The scores are higher with higher cell counts. If there are vast differences in cell type numbers in a dataset to analyze, which was not the case in our study, we recommend dividing COZI scores of a sample by the square root of the number of cells in that sample.

We implemented COZI on a branch of Scimap in order to use the given neighborhood and permutation test framework. We adapted the script to normalize the number of interactions “conditional” and also output the calculated z-score. Therefore, the method follows the SEA approach of calculating a z-score with another interaction count averaging. We ran the method with a Delaunay graph neighborhood definition for the simulated and knn (k=5) for the biological dataset and permuted the cell type labels *n* = 300 times. We obtained a sample-by-z-score matrix as output for downstream evaluation.

### Method comparison

#### Systematic comparison with simulated data

Our basic rationale for systematic method comparison was to compare the abilities to distinguish between different cohorts based on tissue architecture. Therefore, we compared between different cell-cell adjacency groups (random, weak, strong) with similar cell type abundances.

We quantitatively compared the NEP methods on simulated data by determining how well cohorts were classified and how tissue architecture was recovered. The measured features were cell-cell adjacencies (e.g. 0-0, 0-1, …). We trained a random forest model (*caret* package in R) per cohort comparison with the feature vectors per sample. We did not aim to find the ideal model to perform cohort distinction but to identify classification performance differences between the methods. We performed an 80/20 train-validation split with 5-fold cross-validation.

We applied the F1-score as a quantitative comparison measure. The F1-score is the harmonic mean between precision and recall (Formula 2).

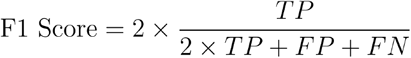

Where:

*TP* = true positive values of classification

*FP* = false positive values of classification

*FN* = false negative values of classification

An F1-score of 1 is the best possible result, while in a binary classification, an F1-score of 0.5 is random. We also generated a baseline for classifying the cohorts based on cell type abundances. And we added a random model where we randomly shuffled the cohort labels.

We evaluated whether the true tissue features were used for cohort distinction by assessing the cosine similarity between the ground truth and the result vectors (Formula 3).

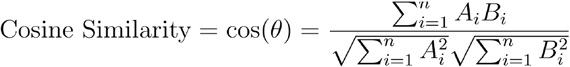

Where:

*A*_*i*_ = *i-*th components of ground truth vector *A* of NEPs

*B*_*i*_ = *i-*th components of NEP result vector *B* of NEPs

The ground-truth vector *A* was created by vectorizing the cell-cell adjacency matrices (Tables 3 and 4). We subtracted the two ground-truth cell-cell adjacency vectors that were compared to generate the ground-truth comparison vector. The result vector *B* included the extracted feature importances of the comparison-specific trained random forest model. The cosine similarity was scaled to a scale between 0 and 1, 1 being perfect similarity and 0 being dissimilarity (Formula 4).

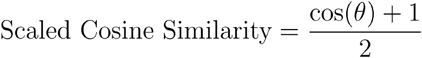

For Giotto, the only tool providing one score per neighbor preference pair, we copied the given scores for an NEP A-B to represent B-A. Per comparison, we determined 100 F1 and 100 cosine similarity scores and plotted them per cell-cell adjacency and cell-type abundance group per tool. Both scores can be compared across methods.

#### Performance comparison on biological data

We ran all described NEP methods on the mouse myocardial infarction dataset by Wuennemann *et al*.^25^ studying immune cell infiltration into the infarct core through the endocardial layer. We used the provided cell type labels for NEP analysis, excluding cell types with label “exclude”. We used a k=5 k-nearest neighbor neighborhood definition. Otherwise, we used the Delaunay neighborhood definition.

We showed cell type abundances in specific regions in the manuscript. We selected the infarct core and the two border zones for the region “infarct region” and the region Endocardial layer for the region “Endocardial infarct zone”.

## Data availability

IST data can be reproduced with the provided scripts in this repository: https://github.com/SchapiroLabor/IST_generation_SCNA

Sequential Immunofluorescence is available via Synapse (project SynID: syn51449054): https://www.synapse.org/Synapse:syn51449054.

The dataframe with phenotypes is available via Synapse: https://www.synapse.org/Synapse:syn65487454.

## Code availability

The github repository for IST generation is:

- https://github.com/SchapiroLabor/NEP_IST_generation

The fork of Scimap where COZI ist implemented

- https://github.com/chiarasch/scimap

The github repositories for running all NEP methods are:

- https://github.com/SchapiroLabor/NEP_Giotto (Giotto)
- https://github.com/SchapiroLabor/NEP_IMCRtools (IMCRtools classic and HistoCAT
- https://github.com/SchapiroLabor/NEP_MistyR (Misty)
- https://github.com/SchapiroLabor/NEP_Squidpy (Squidpy, CellCharter, neighbor counts and cell type abundances)
- https://github.com/SchapiroLabor/NEP_scimap (COZI, SEA and Scimap)

The github repository for systematic tool comparison is:

- https://github.com/SchapiroLabor/NEP_comparison

All repositories contain scripts to reproduce the presented results in this study. IST computations were performed using the Baden-Wurttemberg High Performance Cluster bwForCluster Helix.

## Acknowledgements

The authors gratefully acknowledge the data storage service SDS@hd supported by the Ministry of Science, Research and the Arts Baden-Württemberg (MWK), support by the state of Baden-Württemberg through bwHPC and the German Research Foundation (DFG) through grant INST 35/1314-1 FUGG, INST 35/1503-1 FUGG and INST 35/1597-1 FUGG. We thank Ethan Baker for his input on the *in silico* tissue generation. We are very grateful to Leeat Keren for sharing her code for SEA. We thank Leonie Küchenhoff, Gesa Voigt and Lukas Hatscher for their input throughout the study and Victor Perez for revising the final manuscript. We acknowledge the use of OpenAI’s ChatGPT in this project which assisted by adapting the language to improve clarity and coherence while ensuring that the original scientific content remained intact.

C.S. is supported by the Bruno and Helene Jöster Stiftung. C.S., M.I. and K.B. are supported by the German Federal Ministry of Education and Research (BMBF 01ZZ2004). J.T. is supported by the Ministry for Science, Research and Science Baden-Württemberg “MULTI-SPACE”, the Bruno and Helene Jöster Stiftung and the Multi-dimensionAI project (CZS-Project number: P2022-08-101) was made possible by funding from the Carl-Zeiss-Stiftung. D.S. is supported by the German Federal Ministry of Education and Research (BMBF 01ZZ2004); the Ministry for Science, Research and Science Baden-Württemberg „AI Health Innovation Cluster” and “MULTI-SPACE”; research funding from Cellzome, a GSK company, the Bruno and Helene Jöster Stiftung and the Multi-dimensionAI project (CZS-Project number: P2022-08-101) was made possible by funding from the Carl-Zeiss-Stiftung.

## Author information

### Contributions

C.S., M.I. and D.S. conceived and designed the study. C.S. simulated the data, implemented the methods, designed and performed the method comparison. M.I., J.T. and D.S. supervised the study. M.I and C.S. performed Misty analysis. K.B. and C.S. created plots and analysis and K.B. provided input for the biological MI dataset. C.S. and D.S. wrote the manuscript, revised by M.I., K.B. and J.T. All authors read, discussed and approved the final manuscript.

