## Supplementary Figures 1-9 for "Comparison and Optimization of Cellular Neighbor Preference Methods for Quantitative Tissue Analysis"

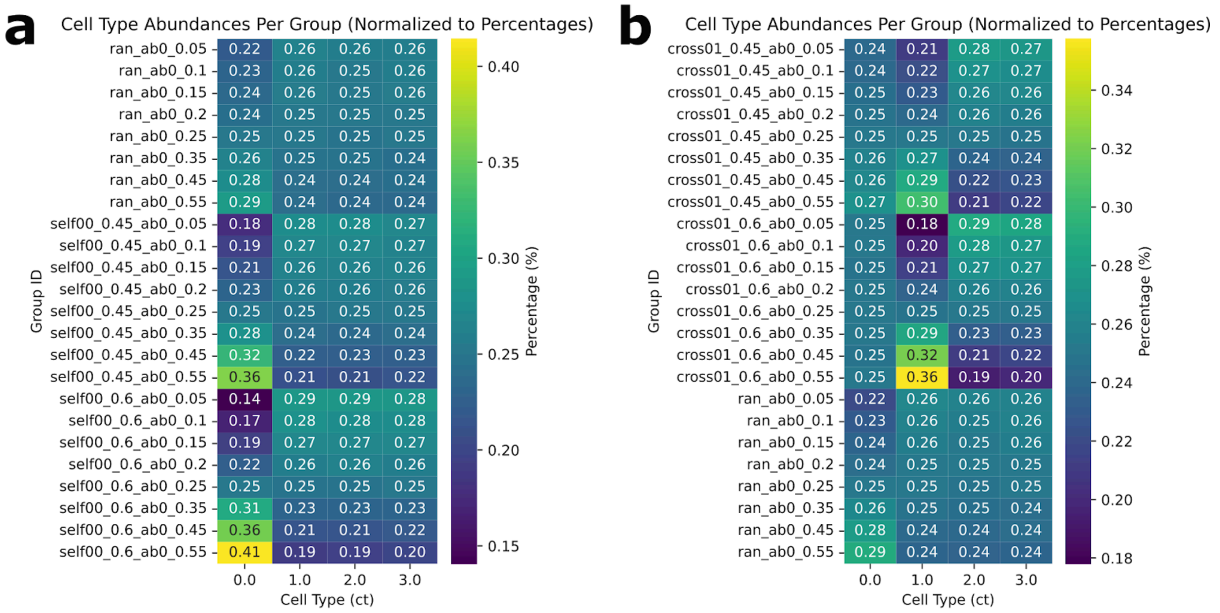


**Supplementary Fig. 1: Differences in cell type abundances between cohorts (a)** Symmetric self-preference of ct 0 (red cell type). **(b)** Asymmetric cross-preference of cell type 0 to cell type 1 (red to yellow). Percentages of cell type abundances per simulated abundance and adjacency cohort. In each cohort, the abundance of ct 0 was set to the respective abundances indicated in the rownames (see Methods).

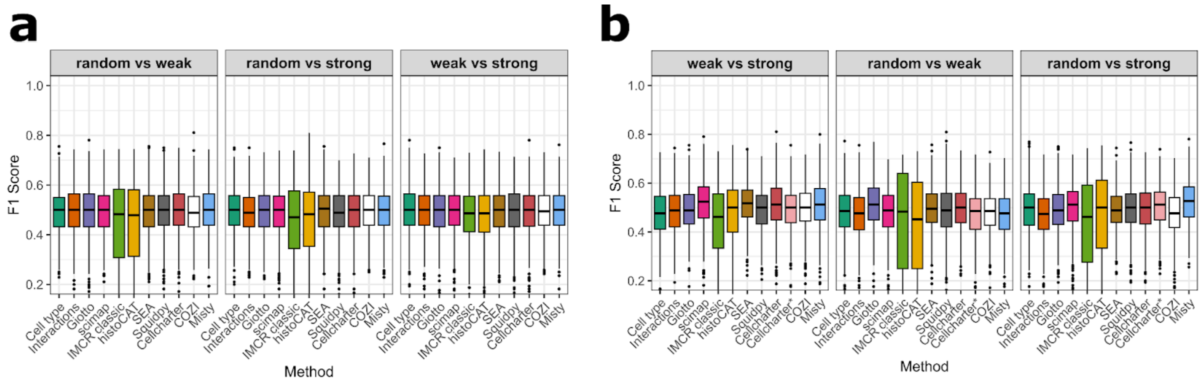


**Supplementary Fig. 2: Baseline F1 scores after random label shuffling of NEP results.** **(a)** Baseline mean F1-scores per cohort distinction task of self-preference of cell type 0 dataset across yellow cell type abundance groups are shown as boxplots. Cohort comparisons were weak vs. strong, random vs. weak and random vs. strong while cohort labels were randomly shuffled. **(b)** Baseline mean F1-scores per cohort distinction task of cross-preference of cell type 0 to 1 dataset across yellow cell type abundance groups are shown as boxplots. Cohort comparisons were weak vs. strong, random vs. weak and random vs. strong while cohort labels were randomly shuffled.

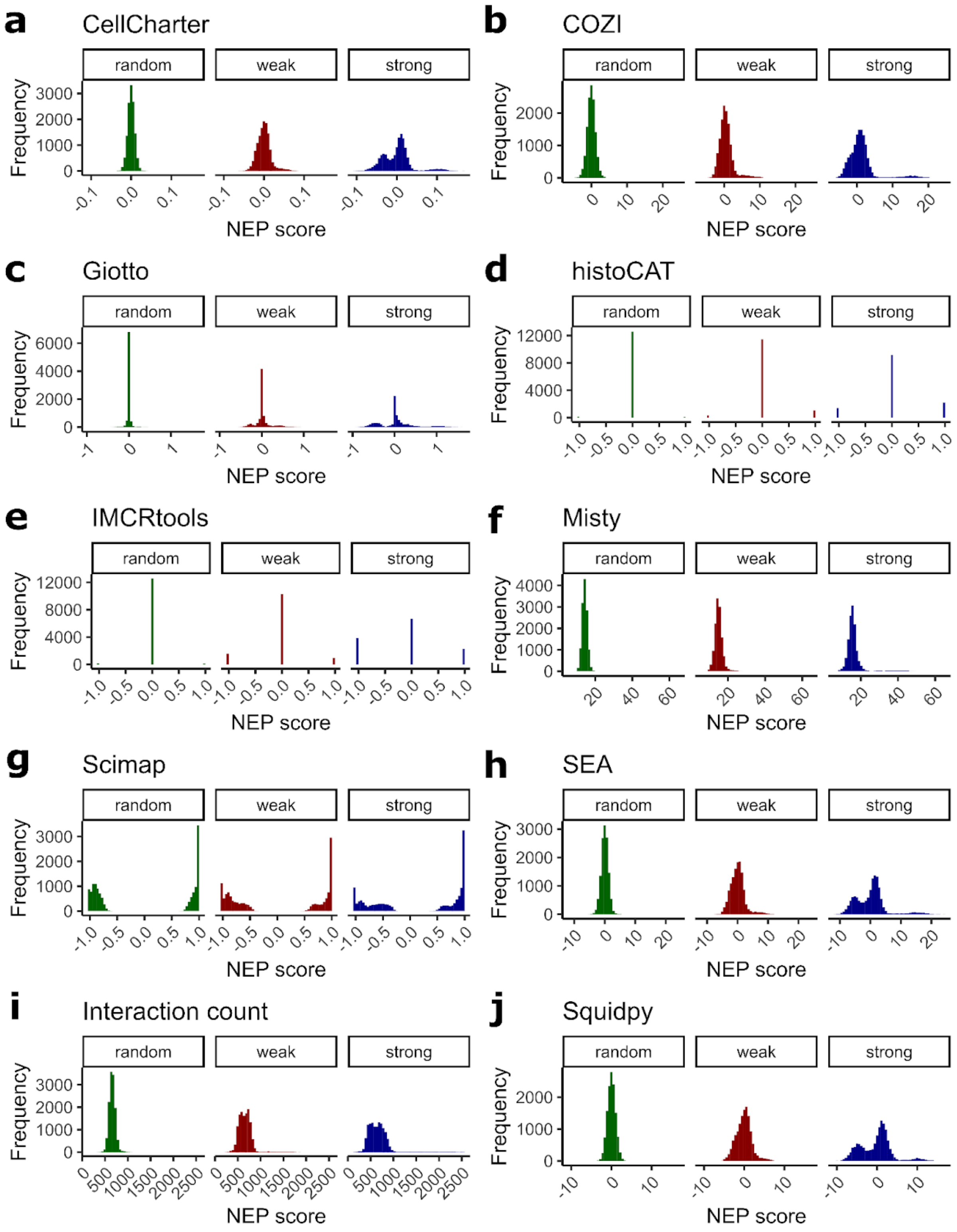


**Supplementary Fig. 3: Overall NEP score frequencies across methods in simulated symmetric tissue cohorts. (a)-(j)** Overall NEP score distributions per method across all cell type pairs in random (green), weak (red) and strong (blue) simulated tissue cohorts. Scores across all abundance groups are depicted.

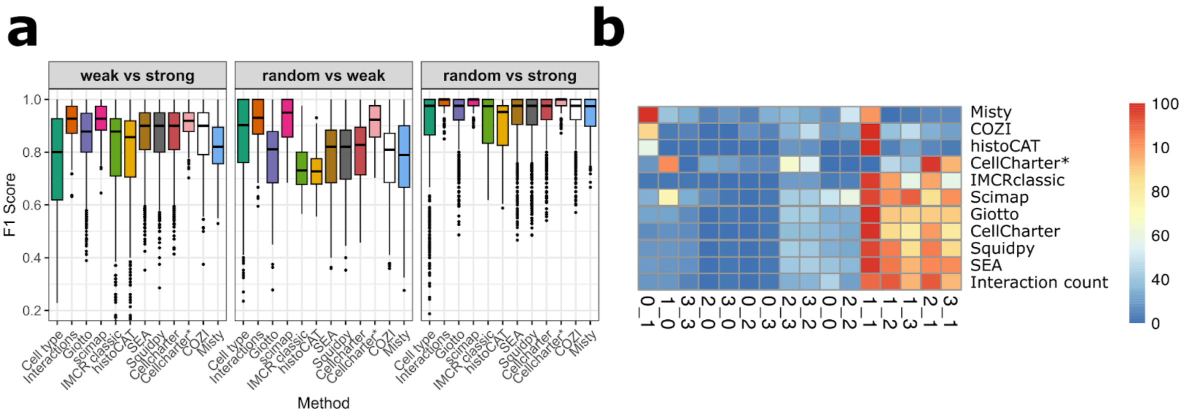


**Supplementary Fig. 4: Overall cohort classification of cross-preference dataset. (a)** Mean F1-scores per cohort distinction task of cross-preference of cell type 0 to 1 dataset across yellow cell type abundance groups are shown as boxplots. Cohort comparisons were weak vs. strong, random vs. weak and random vs. strong. **(b)** Feature importances of the cohort distinction between random and strong self-preference cohorts with a cell type 0 and 1 abundance of 25%.

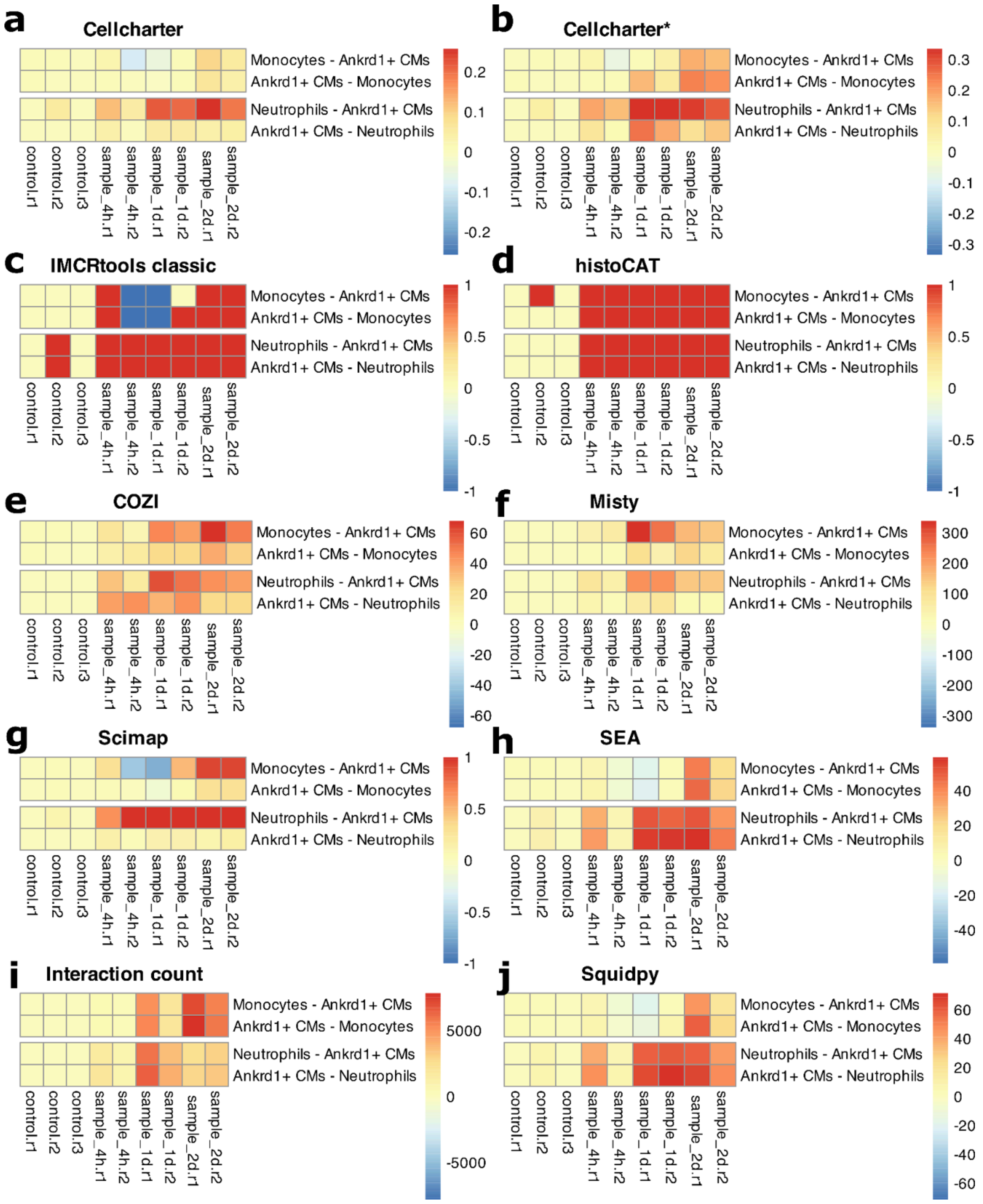


**Supplementary Fig. 5: NEP scores of compared methods for monocyte and neutrophil infiltration into the infarct region.** NEP scores for monocytes and neutrophils with Ankrd1+ cells in control, 4, 24 and 48h samples. **(a)-(h)** CellCharter, HistoCAT, IMCRtools classic, COZI, Misty, Scimap, SEA, Interaction count and Squidpy scores are shown.

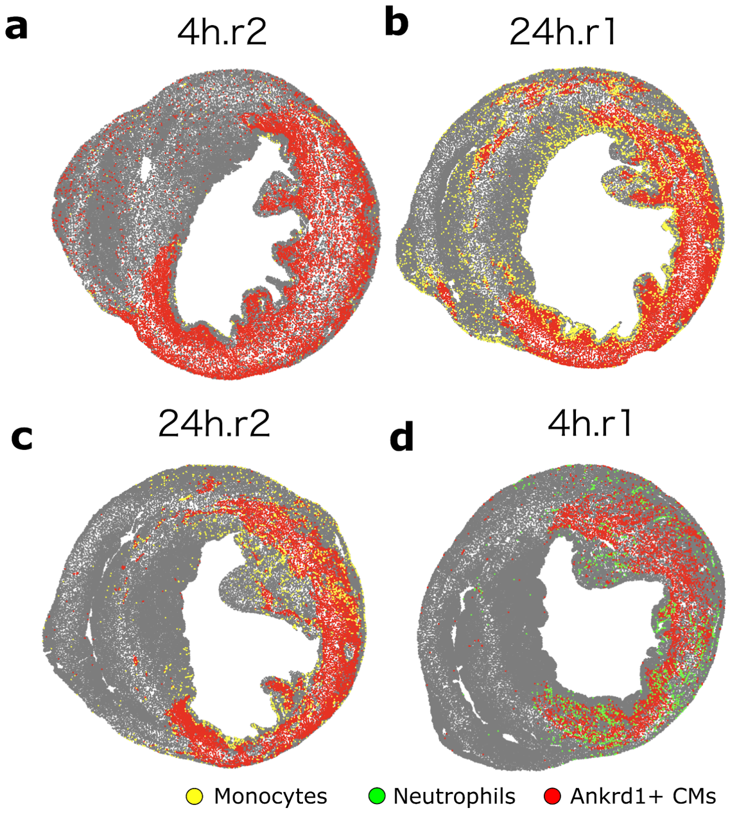


**Supplementary Fig. 6: Mouse hearts at 4 and 24 h after infiltration to study monocyte and neutrophil infiltration.** Samples at 4 h for monocyte **(a)** and neutophil **(d)** infiltration, Both samples at 24h for monocyte infiltration **(b, c)**. Monocytes (yellow), neutrophils (green) and Ankrd1+ cardiomyocytes (red) as scatter plots.

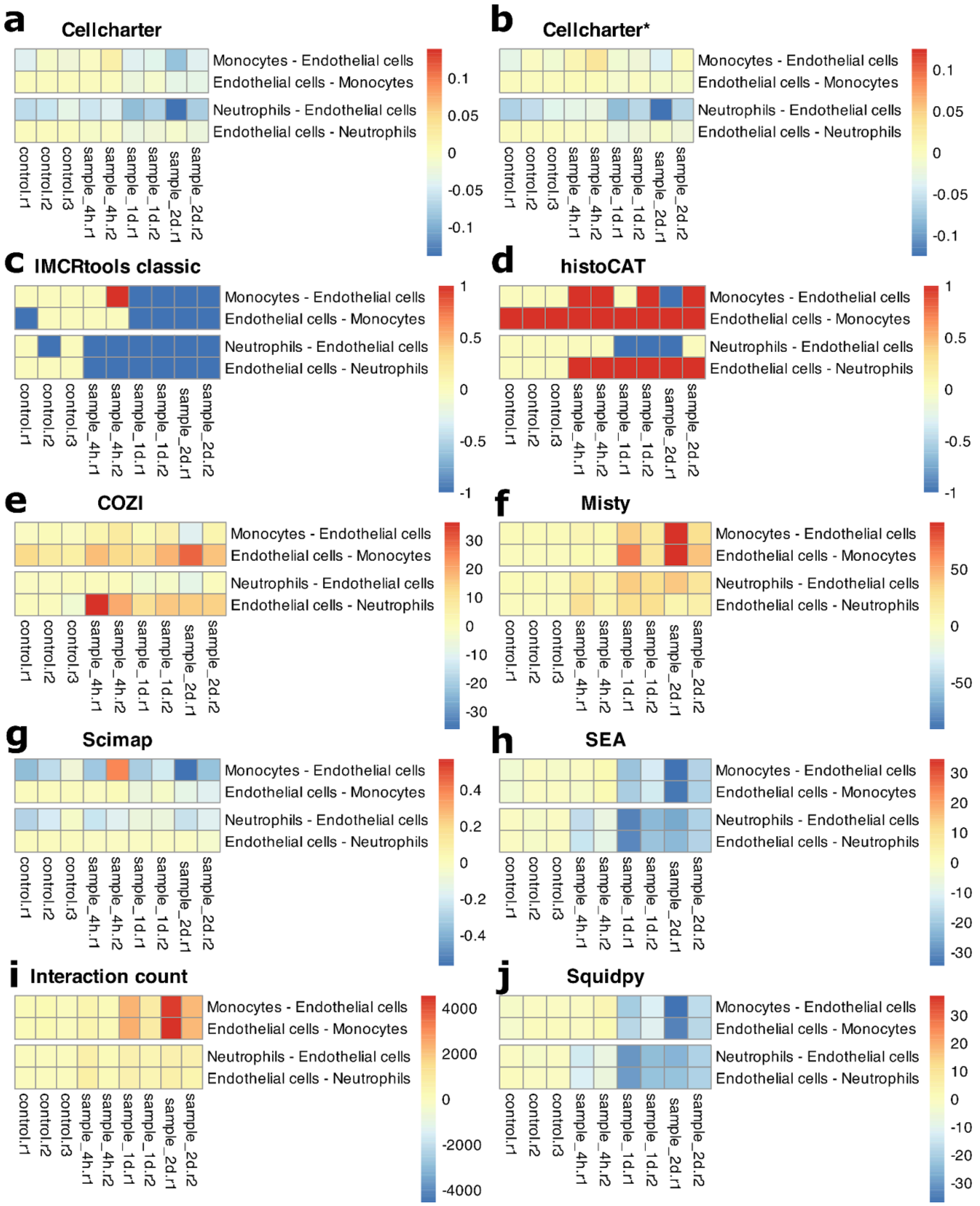


**Supplementary Fig. 7: NEP scores of compared methods between monocytes and neutrophils with endothelial cells.** NEP scores for monocytes and neutrophils with Endothelial cells in control, 4, 24 and 48h samples. **(a)-(h)** CellCharter, HistoCAT, IMCRtools classic, COZI, Misty, Scimap, SEA, Interaction count and Squidpy scores are shown. **(i)** Monocyte infiltration through blood vessels. DAPI (blue), MPO (magenta) and CD31 (yellow) mark cell nuclei, neutrophils and endothelial cells, respectively. The scale bar indicates 50 microns. **(j)** Cell type counts across samples for monocytes, neutrophils, Ankrd1+ cardiomyocytes and endocardial cells.

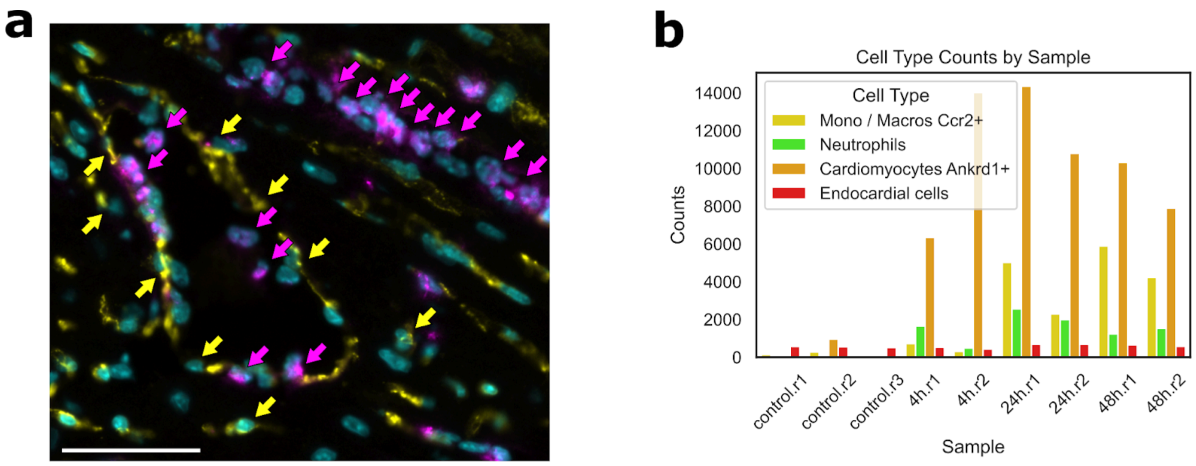


**Supplementary Fig. 8: Neutrophil infiltration through the endocard. (a)** Neutrophil infiltration through blood vessels. DAPI (blue), MPO (magenta) and CD31 (yellow) mark cell nuclei, neutrophils and endothelial cells, respectively. The scale bar indicates 50 microns. **(b)** Cell type counts across samples for monocytes, neutrophils, Ankrd1+ cardiomyocytes and endocardial cells.

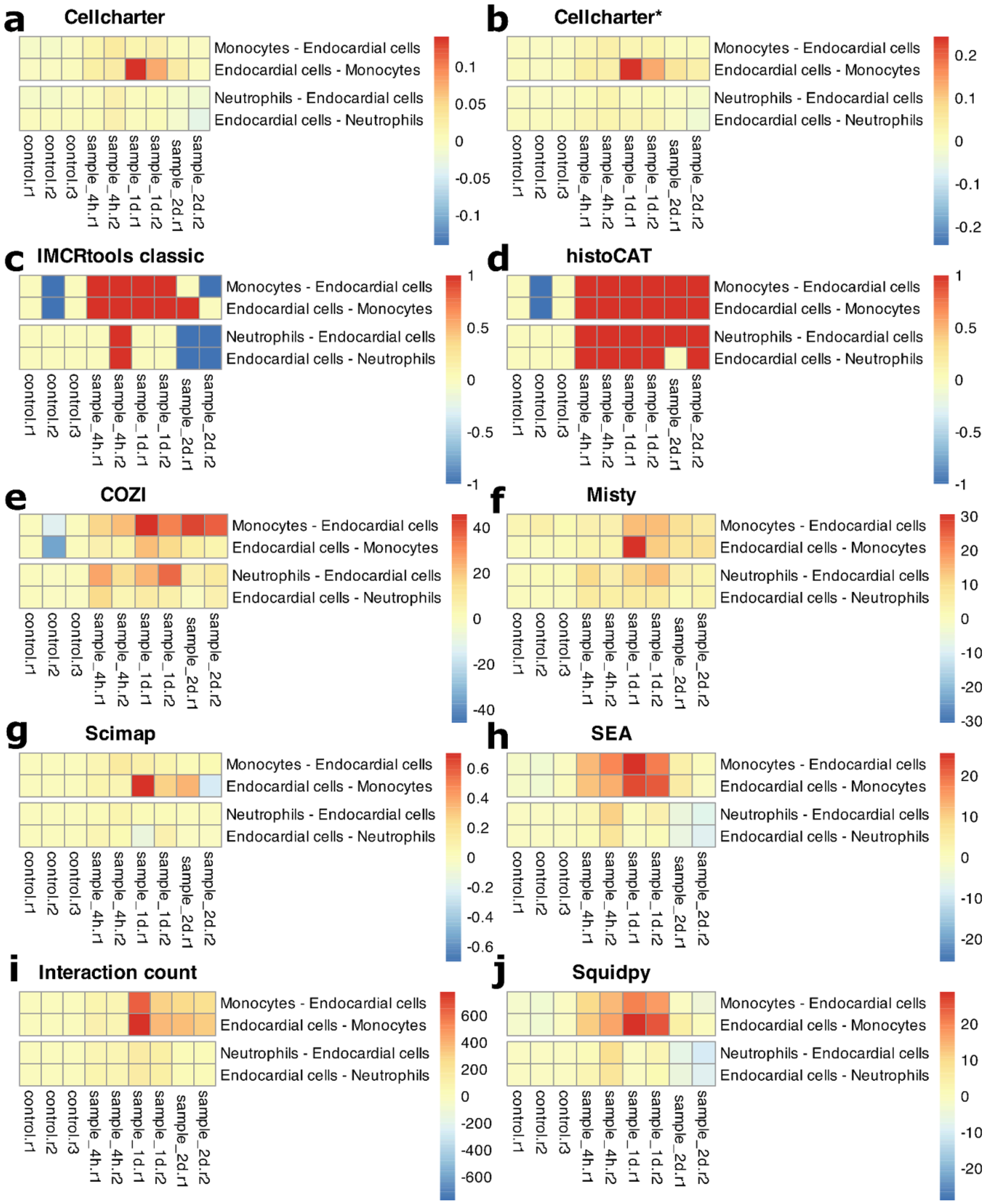


**Supplementary Fig. 9: NEP scores of compared methods between monocytes and neutrophils with endocardial cells.** NEP scores for monocytes and neutrophils with Endothelial cells in control, 4, 24 and 48h samples. **(a)-(h)** CellCharter, HistoCAT, IMCRtools classic, COZI, Misty, Scimap, SEA, Interaction count and Squidpy scores are shown.
